# Proliferation symmetry breaking in growing tissues

**DOI:** 10.1101/2024.09.03.610990

**Authors:** Xinzhi Li, Aniruddha Datta, Shiladitya Banerjee

## Abstract

Morphogenesis of developing tissues results from anisotropic growth, typically driven by polarized patterns of gene expression. Here we propose an alternative model of anisotropic growth driven by self-organized feed-back between cell polarity, mechanical pressure, and cell division rates. Specifically, cell polarity alignment can induce spontaneous symmetry breaking in proliferation, resulting from the anisotropic distribution of mechanical pressure in the tissue. We show that proliferation anisotropy can be controlled by cellular elasticity, motility and contact inhibition, thereby elucidating the design principles for anisotropic morphogenesis.

---

Tissue morphogenesis is primarily driven by anisotropic growth [1–9], leading to the formation of directionally biased organ shapes. For instance, during *Drosophila* gastrulation, polarized intercalations, cell divisions, and motility work together to narrow the extending germband along the dorsalventral axis while elongating it more than twice its length along the anterior-posterior axis [2]. The development of *Drosophila* imaginal wing discs is another striking example of anisotropic morphogenesis, where polarized cell divisions shape the anisotropic organ [3, 4, 10–12]. Anisotropic growth is central to determining shapes in various organisms, including jaw-joint morphogenesis in zebrafish [13, 14], limb bud development in vertebrates [15, 16], and plant morphogenesis [17]. A key question is how cells collectively achieve a directional bias to guide this anisotropic growth.

One prevailing model is that tissue growth and morphogenesis are regulated by biochemical signaling, through patterned gene expression [18]. However, studies over the past decade have highlighted the significant role of mechanical forces in shaping tissue growth, size, and form [1, 19–21]. Mechanical forces can influence local tissue growth rates [22–25], the rate and orientation of cell division [26–29], while cell growth and proliferation can, in turn, affect tissue mechanics through the generation and dissipation of active stresses [30, 31]. This interplay between growth and mechanics suggests a complex crosstalk, though their specific roles in regulating anisotropic growth are not yet fully understood. In this study, we demonstrate that feedback between tissue growth and mechanics can spontaneously break proliferation symmetry to generate anisotropic growth.

We developed a cell-based model for anisotropic tissue growth that integrates feedback between cell motility, mechanics, and cell cycle dynamics by extending the framework of cellular vertex models. Our model implements cell polarity dynamics and its coupling with tissue mechanics, showing that polarized growth can result from the anisotropic distribution of mechanical pressure driven by spontaneous cell polarization. The direction of tissue growth is determined by the polarity axis: in areas with lower compression at the tissue rear, cell divisions occur more frequently, whereas in regions under greater compression at the front, cell divisions are less frequent. Using cell-based modeling and continuum theory, we show that the extent of division anisotropy can be tuned by modulating cell elasticity, motility and contact inhibition, thereby revealing the design principles for anisotropic morphogenesis in growing tissues.

## Discrete cell-based model

We constructed a cell-based model for a growing confluent cell monolayer using the framework of the vertex model [32–38], where the tissue is represented by network of cellular polygons that tessellate the plane. Tissue mechanics is governed by the energy function:

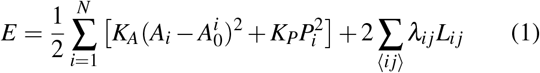

where the cell areas {*A*_*i*_} and perimeters {*P*_*i*_} are functions of the position of vertices {**r**_*i*_} and the connectivity between cells, *K*_*A*_ and *K*_*P*_ are the area and perimeter elastic moduli, and ∑ _⟨*ij*⟩_ represents the sum over all neighboring cells of cell *i*. The term quadratic in cell area *A*_*i*_ results from cell volume incompressibility [33, 34, 36], and 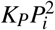 is the elastic energy associated with the actomyosin cortex. The third term describes the line tension, *λ*_*ij*_, along the interface shared by cells *i* and *j* of length *L*_*ij*_, which can be up-regulated by increasing actomyosin contractility or by lowering cell-cell adhesion.

Cell growth is implemented by increasing the cell’s target area 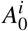 [39, 40] as 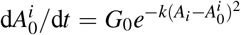, where the constant *G*_0_ is the growth rate of individual cells in the absence of crowding, and *k* is the cellular sensitivity to crowding. In crowded conditions, cells are under compression which causes deviations of cell area from its preferred value in isolation, leading to an increase in cell pressure 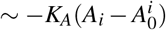. This results in a decrease in cell area growth rate at a rate *k*, thereby lengthening cell cycle times. The parameter *k* thus models the effect of contact inhibition of proliferation [20], through mechanical feedback on growth. The number of cells in the tissue *N* increases through cell divisions, regulated by a G1 sizer model [41–43]. In this model, a cell grows in the G1 phase until its area is greater than a threshold value *A*_*S*_, at which point it transitions to the S/G2/M phase where it resides for a fixed time period *T* (a timer) before dividing into two daughter cells (see Supplemental Material).

In addition to growth and division, each cell can migrate actively. To model active cell migration, each cell is assigned a polarity vector ***p***_*i*_ that defines the direction of migration. The overdamped equation of motion of cell vertex *α* is:

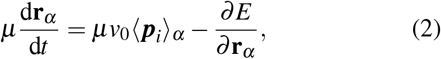

where *µ* is the friction coefficient with the substrate, *v*_0_ is the cell crawling speed, and ⟨***p***_*i*_⟩_*α*_ is the average polarity vector of cells sharing the vertex *α*. With the given rules for polarity dynamics (discussed below), we simulated the vertex-based model of tissue growth, starting from a small colony of *N*_0_ cells under free boundary conditions (Fig. S1). See Supplemental Material for details on simulation methods and parameter choices. When a cell reaches the timer threshold, it divides by splitting perpendicular to its major axis, with the boundaries of the daughter cells passing through the center of the mother cell (Fig. S1). The preferred areas of the daughter cells are determined by maintaining pressure homeostasis within cells before and after division. Following division, the ages of the daughter cells are reset to zero, and their polarity vectors are randomly assigned. The network topology is updated using T1 transitions to facilitate neighbor exchanges.

### Isotropic growth

In the absence of any directional bias, the polarity vector of each cell undergoes rotational diffusion, resulting in isotropic growth (Fig. 1a, Movie 1). Cell divisions occur mostly at the tissue boundary, recapitulating experimental observations [28, 29]. This is due to crowding-induced suppression of growth at the bulk of the tissue, where the cells undergo cell cycle arrest (Fig. S1). As a consequence, there is a buildup of mechanical pressure at the center of the colony (Fig. 1b, Movie 2) and cell growth rate decreases from the boundary to the center of the tissue (Fig. S1). Importantly, there is a negative correlation between cell pressure and the probability of cells being in the G2 phase, such that cell division is more likely in regions of low pressure near the tissue boundary (Fig. 1c, Fig. S2). This highlights the role of mechanical feedback on the patterning of cell proliferation. The emergent patterns of proliferation can be controlled by modulating cell elasticity *K*_*A*_ and the contact inhibition parameter *k* (Fig. S2).

**FIG. 1.**
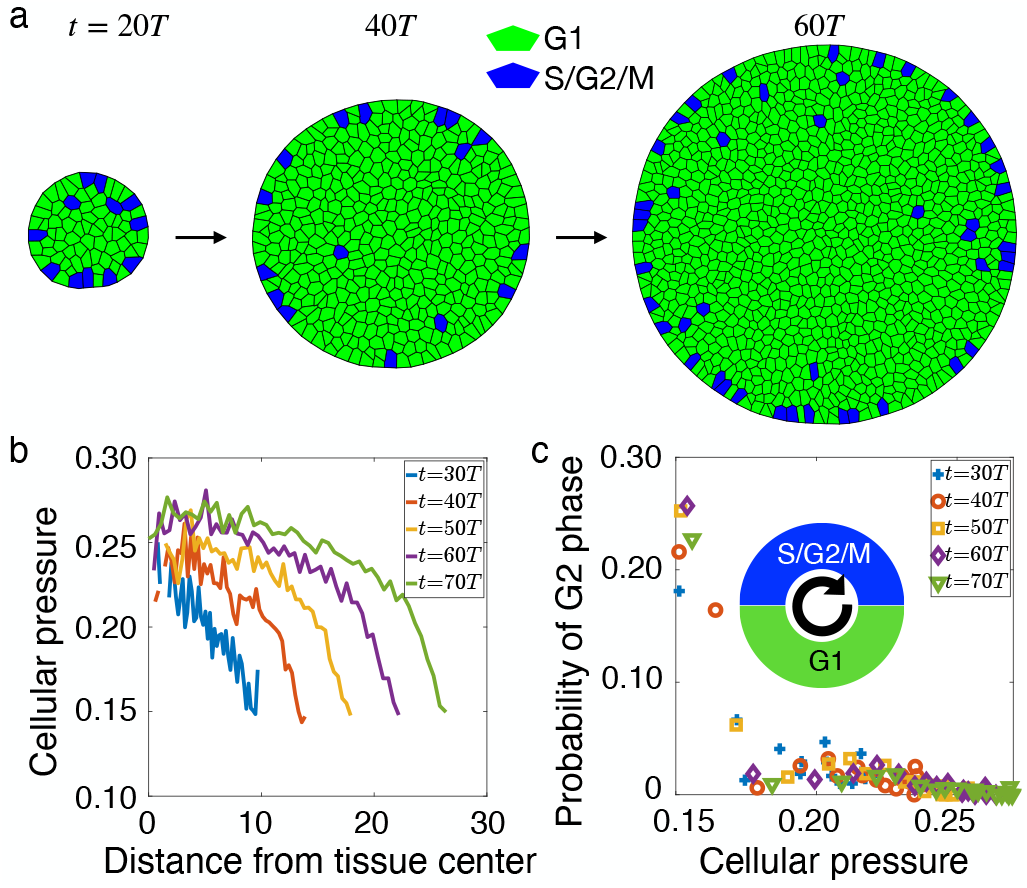
Proliferation pattern in isotropic growth. (a) Timelapse of an isotropically growing tissue with random polarity. The colors represent cells in G1 (green) and G2 (blue) phases of the cell cycle. (b) Radial pressure distribution at various time steps. The crowded inner cells are under higher pressure from neighboring cells which leads to lower growth rate. (c) Correlation between the probability of cells in G2 phase of the cell cycle with the cellular pressure. Inset: Schematic of cell cycle phases. See Table I in the Supplemental Material for a list of parameter values.

### Emergence of anisotropic growth

To more realistically model cell polarity dynamics, we incorporated two essential biological features: contact regulation of locomotion [44] and polarity alignment with neighbors [45, 46]. Due to contact regulation of locomotion, cells on the boundary of the monolayer tend to migrate into free space, setting their initial polarity vectors normal to their free edges and pointing outwards. Cells within the monolayer align their polarity vectors with their neighbors [47]

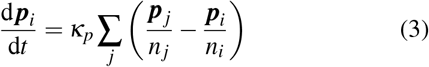

where *j* labels the neighboring cells of the cell *i, κ*_*p*_ is the polarity alignment rate, *n*_*i*_ and *n* _*j*_ are the number of neighbors for cells *i* and *j*, respectively. With these rules, the growing tissue develops an anisotropic distribution of cell growth rate and polarity (Fig. 2a-b), anisotropy in division orientation (Fig. 2c) and irregular shape due to differential growth (Movies 3, 4).

**FIG. 2.**
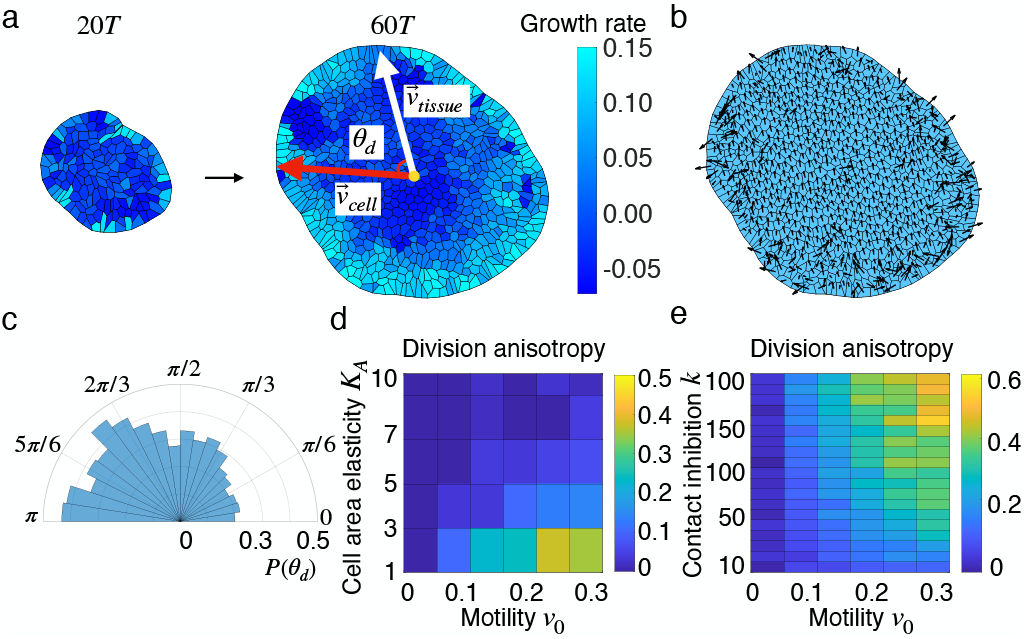
Anisotropic division patterns in polarized tissues. (a) Timelapse of an anisotropically growing tissue. Colormap represents cell growth rate which shows anisotropic distribution within the tissue. On the right panel, we schematically depict the calculation of cell division angle 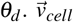 is the vector pointing from the tissue center to the center of the dividing cell and 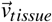 represents the tissue axis defined as the average of all cellular polarity vectors within the tissue. (b) Cell polarity distribution. (c) Anisotropic angular distribution of cell divisions. Fewer cell divisions in the head part of the tissue with 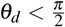 while more in the rear part with 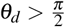. (d) Phase diagram of division anisotropy at various values of cell area elasticity *K*_*A*_ and migration speed *v*_0_. (e) Phase diagram of division anisotropy at various values of the contact inhibition of proliferation *k* and *v*_0_. See Table I in the Supplemental Material for a list of parameter values.

To characterize the anisotropy of cell divisions, we calculated the angles of cell division θ_*d*_, defined as the angle between the mean polarity vector and the vector joining the tissue center to the center of the dividing cell (Fig. 2a-right). 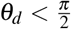 indicates cell divisions in the head part of the tissue, while that in the rear part implies division angles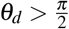. The polar histogram in Fig. 2c shows the anisotropic distribution of cell divisions, with the head part exhibiting less divisions compared to the rear. We introduce the division anisotropy factor, *d*_ans_ = 1− *N*_*h*_/*N*_*r*_, as a metric of anisotropic growth, where 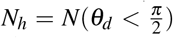 and 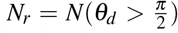 are the numbers of division events in the head and rear parts, respectively. When cells divide isotropically, *d*_ans_ = 0, while *d*_ans_ = 1 indicates that all divisions occur in the tissue rear. To investigate how tissue properties control division anisotropy, we calculated *d*_ans_ at various cell area elasticity *K*_*A*_ and cell motility *v*_0_ values. The phase diagram in Fig. 2d shows that increasing *v*_0_ increases *d*_ans_, while increasing *K*_*A*_ decreases *d*_ans_. This is understandable since directional motility (*v*_0_) makes polarity alignment more efficient while *K*_*A*_ imposes stronger constraints on cell area and enhances uniformity. The contact inhibition parameter *k* also plays a key role in regulating division anisotropy, such that with the increasing *k*, cells become more sensitive to tissue crowding and thus the division anisotropy is enhanced (Fig. 2e). In the absence of mechanical feedback on growth (*k* = 0), division anisotropy is lost (Fig. S3). Overall, changing *k, v*_0_ and *K*_*A*_ modulates division anisotropy, and also influences tissue morphology (Figs. S4-S5). While it is expected that polarity alignment would induce a flocking transition to drive directional collective motion [48], the role of cell polarity in coordinating anisotropic divisions is novel, which we next address in detail.

### Mechanical origin of proliferation symmetry breaking

To uncover the mechanical origin of division anisotropy, we analyzed cell pressure distribution, as pressure negatively correlates with the probability of cell division (Fig. 1c). We compared pressure distributions in a growing isotropic tissue, a growing anisotropic tissue, and a tissue without cell proliferation (Fig. 3a). In the absence of polarity alignment (Fig. 3a, left) or mechanical feedback on growth (*k* = 0, Fig. S3), the tissue grows isotropically with an isotropic pressure distribution that decreases from the center to the boundary. When polarity alignment is present, the pressure distribution is anisotropic, corresponding to anisotropic growth (Fig. 3a- middle, Movie 5). Here, cell divisions are less frequent in regions of high pressure, and more frequent in low-pressure areas. Additionally, simulations of a non-proliferating tissue revealed that polarity alignment alone is sufficient to induce anisotropic pressure oriented along the polarity axis (Fig. 3a, right), suggesting that pressure anisotropy is a precursor to division symmetry breaking. Quantification of angular pressure distribution (Fig. 3b) confirms that polarity alignment is indeed essential for anisotropic pressure.

Collectively, these results demonstrate that the symmetry breaking in proliferation is driven by the emergent anisotropy in pressure, which in turn is induced by a gradient in cell compression from the head to the rear of the tissue. This gradient results from tissue deformation induced by cell polarization, suggesting that cell elasticity can regulate the degree of pressure anisotropy. To quantify this, we calculated pressure anisotropy as *p*_*ans*_ = 1 − *p*_*r*_/*p*_*h*_, where *p*_*r*_ and *p*_*h*_ are the average pressures in the rear (*θ* > *π*/2) and the head (*θ* < *π*/2) regions of the tissue, with *θ* representing the angle of each cell relative to the tissue center. We found that increasing cell area elasticity (*K*_*A*_) suppresses pressure anisotropy (Fig. 3c), while increasing cell motility (*v*_0_) and contact inhibition (*k*) enhances it (Fig. 3d). Furthermore, pressure anisotropy is negligible in the absence of contact inhibition of proliferation (Fig. 3d, Fig. S3). These trends align with that of cell division anisotropy in Fig. 2d-e, suggesting that pressure anisotropy underlies asymmetric proliferation patterns.

**FIG. 3.**
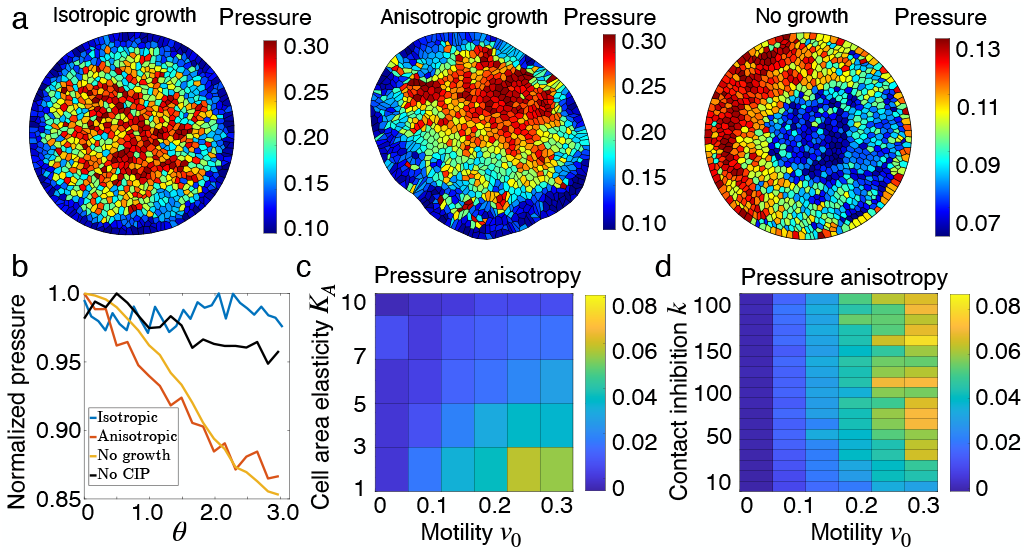
Pressure anisotropy underlies proliferation symmetry breaking. (a) Snapshots of pressure distribution for an isotropically growing tissue (left), anisotropically growing tissue with polarity alignment (middle) and a tissue system without cell proliferation but under polarity alignment (right). The colors represent magnitudes of cell pressures. (b) Quantification of pressure anisotropy for the tissue configurations in (a) and the case without contact inhibition *k* = 0. For each configuration, the pressure is normalized by its maximum value. (c) Phase diagram of pressure anisotropy at various values of cell area elasticity *K*_*A*_ and migration speed *v*_0_. (d) Phase diagram of pressure anisotropy at various values of contact inhibition *k* and *v*_0_.

### Continuum theory

To gain physical insights into the emergence of anisotropic growth, we formulated a minimal continuum model of a growing tissue. With the continuum model we show that our proposed mechanism for proliferation symmetry breaking does not depend on the specific microscopic rules of the vertex model, but is a collective phenomenon resulting from the feedback between cell polarity, mechanical pressure, and proliferation. It suffices to illustrate the underlying physics using a one-dimensional model of a growing tissue, characterized by three time-varying fields, namely density *ρ*(*x, t*), velocity *v*(*x, t*) and polarity *p*(*x, t*), governed by the following set of equations:

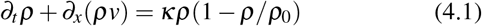

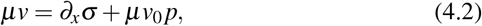

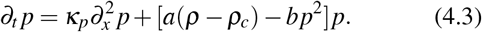

Mass balance implies that the density satisfies the continuity equation (4.1) with logistic growth, where *κ* is the growth rate constant, and *ρ*_0_ is the homeostatic density. The effective rate of exponential proliferation, *κ*(1 − *ρ*/*ρ*_0_), decreases with density, capturing the effect of contact inhibition of proliferation. Eq. (4.2) represents force balance in the overdamped limit, where *σ* is the tissue stress field, *µ* is the friction with the substrate, and *v*_0_ is the speed of active cell crawling. We assume the constitutive equation of Maxwell viscoelasticity since the tissue is expected to behave like a viscous fluid at long times,

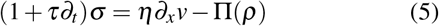

where *η* is the tissue viscosity and Π is the pressure that is dependent on cell density. Lacking precise knowledge of the tissue’s equation of state, we use a first-order expansion of the pressure around the homeostatic density *ρ*_0_, yielding Π(*ρ*) = Π_0_ + *χ*^−1^(*ρ*− *ρ*_0_), where Π_0_ is the homeostatic pressure and *χ* is the compressibility (inverse bulk modulus).

Eq. (4.3) describes cell polarity dynamics, with the first term describing polarity alignment at a rate *κ*_*p*_, and the second term drives symmetry breaking in the homogeneous steady-state when the density is above a threshold *ρ*_*c*_, given *a, b* > 0 (see Supplemental Material).

We analyzed Eqs. (4.1)-(4.3) both analytically and numerically in a domain with no-flux boundary conditions (see Supplemental Material). In the limit *v*_0_ = 0 and *η* = 0, the model reduces to the porous-Fisher model [49], which would exhibit isotropic growth. In the absence of polarity symmetry breaking and polarity alignment (*a* = 0, *κ*_*p*_ = 0), we get isotropic growth with elevated proliferation rate at the tissue boundary (Fig. 4a, Fig. S6). By contrast, a polarized tissue with non-zero *κ*_*p*_ exhibits anisotropic proliferation (Fig. 4b-c), with an overall motion along the direction of polarity. The nonzero polarity field leads to an asymmetry in the velocity field (Fig. S7), which in turn induces an anisotropic distribution of density (Fig. 4b). Growth rate, given by *κρ*(*ρ* − *ρ*_0_), exhibits an anisotropic distribution (Fig. 4c) that captures the spatiallyvarying rate of density change.

**FIG. 4.**
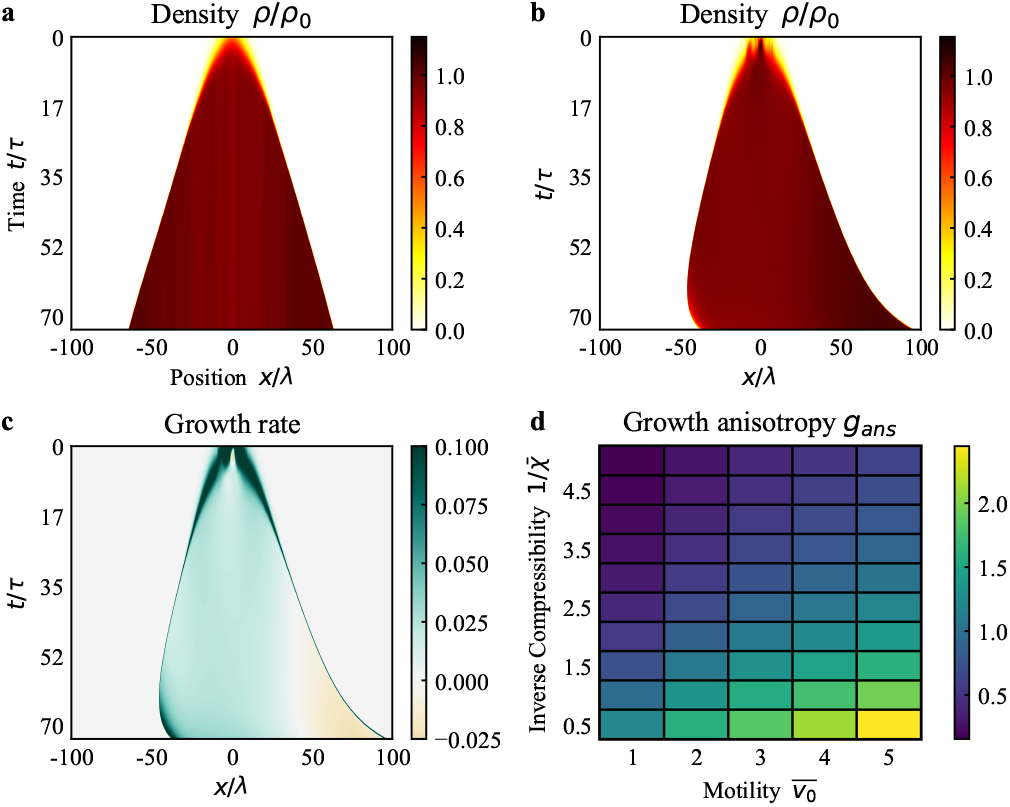
Anisotropic tissue growth in the continuum model. (a) Kymograph of density of an isotropically growing tissue with *a* = 0 and *κ*_*p*_ = 0. (b) Kymograph of density for an anisotropically growing tissue with *a* = 0.3 and *κ*_*p*_ = 1000. The density is higher in +*x* direction compared to −*x* direction. (c) Kymograph of growth rate for the tissue in (b). The growth rate is higher in the negative *x* direction where the density is lower. (d) Phase diagram of growth anisotropy at various values of inverse compressibility 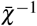 and motility 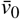. Increasing 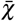 and 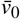 promote growth anisotropy. See Table II in the Supplemental Material for a list of parameter values.

In order to quantify the anisotropy in growth, we defined the growth anisotropy index as *g*_*ans*_ = 1 −*G*_+_/*G*_−_, where *G*_+_ is the sum of the growth rates in the head of the tissue (i.e. the half of the tissue in the direction of the average polarity), and *G*_−_ as the sum of the growth rates in the rear. For an isotropically growing tissue, growth rates are the same in the two halves of the tissue and *g*_*ans*_ = 0 (Fig. S8). For anisotropically growing tissue, the growth rate is lower in the head than in the rear, so *g*_*ans*_ is non-zero. Thus it is analogous to the division anisotropy index *d*_*ans*_ defined in the vertex model. In the continuum model, since the density *ρ* can be greater than *ρ*_0_ locally, the growth rate can be negative and *g*_*ans*_ can exceed 1. We find that *g*_*ans*_ ≈ 0 throughout the simulation for isotropic growth, but increases over time for the anisotropic case after some initial fluctuations (Fig. S8).

To examine how the growth anisotropy is controlled by the model parameters, we calculated *g*_*ans*_ at a fixed tissue size for different values of normalized compressibility 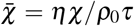 and motility 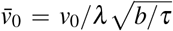, where 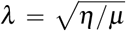. From Fig. 4d, we see that increasing 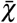 and 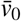 increases the growth anisotropy, mirroring vertex model results (Fig. 3d). To understand this, it is instructive to examine the model behavior at long time and length scales, *t* ≫ *τ* and *x*≫*λ*, when the model reduces to the following nonlinear drift-diffusion equation:

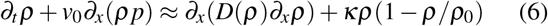

with an effective diffusivity *D*(*ρ*) = *ρ*/(*χµ*). For *v*_0_ = 0, there is no net drift along cell polarity, and the system exhibits isotropic growth. Increasing *v*_0_ amplifies the effect of drift compared to diffusion, driving polarized growth. If compressibility *χ* is high, diffusion is negligible relative to drift. As a result the system maintains density gradients along the direction of cell polarity, thus amplifying growth anisotropy. Taken together, this minimal model establishes the critical role of cell polarity and growth-mechanics feedback in governing anisotropic proliferation.

## Discussion

Anisotropic tissue growth can result from differential proliferation rates [21], directional intercalations [2] or oriented cell divisions set by biochemical gradients [10–12]. Our proposed theory elucidates a new mechanism for self-organized anisotropic growth through feedback between directional cell motility, mechanical stress and cell division. This is relevant for organ morphogenesis driven by active cell migration such as in early limb bud morphogenesis [16, 50, 51], where a coordination of directional cell migration and oriented divisions drives anisotropic organ growth.

## ACKNOWLEDGMENT

We thank Fernanda Perez-Verdugo for many useful discussions. We acknowledge support from the National Institutes of Health (NIH R35 GM143042).

