## Supplemental Material for "Proliferation symmetry breaking in growing tissues"

### I. VERTEX MODEL SIMULATIONS

**Model Initialization and Parameterization**. For the initialization of the vertex model simulation, we generated a set of random cell centers and constructed polygonal cells from voronoi tessellation. The initial state consisted of $N_0$ cells near the center of the system to provide a set of initial vertex positions under free boundary conditions. Each cell is assigned a preferred cell area $A_0^i$ and polarity vector $\boldsymbol{p}_i$ in a random direction with motility speed $v_0$. Edges connected by vertices on the boundary cells act as the constraint from the external environment. The network topology is updated using T1 moves [1, 2] to allow neighbor exchanging and tissue expansion. Parameter values are summarized in Table I.

**Cell growth and division**. In Fig. S1a, we show snapshots of a growing tissue and how cell divisions are implemented in the simulation. Black dots represent cell centers and red dots indicate vertices obtained from voronoi tessellation. As shown in Fig. S1b, when a cell reaches the sizer/timer threshold, it is divided in the perpendicular direction of the main axis (indicated by the red line). The two dark red dots represent the two new vertices connecting the new edge separating the daughter cells which passes through the center of the mother cell. The preferred cell areas of daughter cells are calculated based on the fact that cell pressure remains the same before and after cell division. Assuming the mother cell has area $A$ and preferred area $A_0$, the cellular pressure is $P = -K_A(A - A_0)$. Upon division, the daughter cells obtain area $A_1$ and $A_2$ where $A = A_1 + A_2$. Then the preferred areas of daughter cells are calculated as $A_0^1 = A_1 + P/K_A$ and $A_0^2 = A_2 + P/K_A$. This rule maintains pressure homeostasis within cells. G2 ages of the two daughter cells are reset to zero and the polarity vectors are randomly assigned. A representative single-cell area trajectory is shown in Fig. S1c.

The impact of tissue crowding on cell growth is evident in the single cell area tracks in Fig. S1c. During the early growth phase, cells displayed timer-like behavior, dividing almost at regular intervals of $\sim T$ because their size exceeded the G1 sizer threshold $A_S$. This resulted in size-reductive divisions until the cell size fell below $A_S$, at which point cells grew very slowly in the G1 phase and experienced complete cycle arrest due to their inability to transition to the timer phase. Such patterns of size reductive divisions have been observed experimentally [3]. The emergent patterns of proliferation can be controlled by changing cell elasticity $K_A$ and contact inhibition parameter $k$, as shown in Fig. S2. At a fixed value of the contact inhibition parameter $k$, we observe more cells to be in G2 phase in the bulk by increasing $K_A$. In contrast, increasing $k$ at a given $K_A$ value, cells are more sensitive to crowding and fewer boundary cell divisions could occur. In the absence of contact inhibition, $k = 0$, tissue growth is homogeneous and divisions occur uniformly and isotropically (Fig. S3).

**Cell polarity dynamics**. In simulations of isotropic tissue growth, we suppress polarity alignment interactions. In that case, we model the dynamics of cell polarity vectors as undergoing rotational diffusion [4–6],

$$\begin{aligned} \partial_t \theta_i &= \eta_i(t), \\ \langle \eta_i(t)\eta_j(t') \rangle &= 2D_r \delta(t-t')\delta_{ij}, \end{aligned} \tag{1}$$

where $\theta_i$ is the polarity angle that defines $\boldsymbol{p}_i = (cos\ \theta_i, sin\ \theta_i)$, and $\eta_i(t)$ is a white noise process with zero mean and variance $2D_r$. The value of angular noise $D_r$ determines the memory of stochastic noise in the system, giving rise to a persistence time scale $\tau = 1/D_r$ for the polarization vector $\boldsymbol{p}_i$. To induce anisotropic tissue growth, we implement polarity alignment dynamics that is discussed in the main text.

**Characterization of tissue morphology**. The irregular shape of the anisotropically growing tissue is measured by the ratio $I_u/I_v$, where $I_u$ and $I_v$ are the second moments of area along the major and the minor axes. For a polygon with $n$ vertices $\{x_i, y_i\}$ , numbered in counter-clockwise fashion, the second moment of area is given by [7, 8]

$$\begin{aligned} I_{xx} &= \frac{1}{12}\sum_{i=1}^{n}(x_i y_{i+1} - x_{i+1} y_i)(y_i^2 + y_i y_{i+1} + y_{i+1}^2) \\ I_{yy} &= \frac{1}{12}\sum_{i=1}^{n}(x_i y_{i+1} - x_{i+1} y_i)(x_i^2 + x_i x_{i+1} + x_{i+1}^2) \\ I_{xy} &= \frac{1}{24}\sum_{i=1}^{n}\{(x_i y_{i+1} - x_{i+1} y_i) \ldots. \\ &\quad (x_i y_{i+1} + 2x_i y_i + 2x_{i+1} y_{i+1} + x_{i+1} y_i)\} \end{aligned} \tag{2}$$

The centroidal moments are

$$\begin{aligned} I_{uu} &= I_{xx} - Ay_c^2 \\ I_{vv} &= I_{yy} - Ax_c^2 \\ I_{uv} &= I_{xy} - Ax_c y_c \end{aligned} \tag{3}$$

The principle moments $(I_1, I_2)$ and orientations can be obtained by solving eigenvalues and eigenvectors of the matrix

$$I = \begin{bmatrix} I_{uu} & -I_{uv} \\ -I_{uv} & I_{vv} \end{bmatrix}.$$

Then the shape of the polygon can be measured by the ratio of the principle area moments $A_R = I_1/I_2$. By analogy with single cells, we can also use the shape index of the tissue to measure the shape anisotropy, which is defined as $q = P_t/\sqrt{A_t}$. Here $P_t$ and $A_t$ are the perimeter and area of the tissue, respectively. Lower ratio $A_R$ and higher shape index correspond to more deformed irregular tissue shapes.

By calculating the moment of area and shape index of the tissue polygon formed by boundary vertices, we could characterize the irregularity of tissue shapes. As shown in Fig. S4a and c, at $\kappa_p = 0.05$, the ratio $A_R = I_1/I_2$ decreases while the shape index increases with the increasing of motility $v_0$, indicating tissue shapes become more anisotropic. At $v_0 = 0.1$, the ratio $A_R$ (Fig. S4b) increases while the tissue shape index (Fig. S4d) decreases with the polarity alignment rate $\kappa_p$, representing more isotropic regular tissue shapes. When $\kappa_p$ is increasing, the polarity vectors are more efficiently aligned, leading to collective drifting of the tissue which helps maintain regular isotropic tissue shapes. In the main text, we have discussed the indispensable role of contact inhibition in determining anisotropic growth patterns. Therefore, we show the phase diagram of $A_R = I_1/I_2$ and tissue shape index at various contact inhibition and cell motility values in Fig. S5. Active cell motility promotes anisotropic tissue shapes. In contrast, contact inhibition reinforces the uniformity of tissue growth and maintains regular shapes.

### II. CONTINUUM MODEL

#### A. Model derivation and non-diemnsionalization

Here we describe the continuum model of tissue growth in one spatial dimension. This is relevant for planar tissue growth, with translational invariance along one of the in-plane axes. The tissue is characterized by three time varying fields, namely density $\rho(x,t)$, velocity $\mathbf{v}(x,t)$ and polarity $p(x,t)$. The density equation is derived by assuming mass conservation with logistic growth,

$$\partial_t \rho + \partial_x(\rho v) = \kappa\rho(1 - \rho/\rho_0) \tag{4}$$

where $\kappa$ is the rate of proliferation and $\rho_0$ is the homeostatic density. Condition of force balance in the overdamped limit yields the following equation of motion,

$$\partial_x \sigma = \mu v - \mu v_0 p \tag{5}$$

where $\sigma$ is the stress in the tissue, $\mu$ is the coefficient of friction, and $v_0$ is the speed of active cell motility. Active motility occurs along the direction of local cell polarity $p$. The constitutive equation for tissue stress is assumed to follow that of Maxwell viscoelastic materials since the tissue behaves as a fluid at long times due to cell rearrangements and divisions. We thus have,

$$(1 + \tau\partial_t)\sigma = \eta\partial_x v - \Pi(\rho) = \eta\partial_x v - (\Pi_0 + \chi^{-1}(\rho - \rho_0)) \tag{6}$$

where $\tau$ is the timescale of viscoelastic relaxation $\eta$ is the viscosity and $\Pi(\rho)$ is the density-dependent pressure. We assume the pressure $\Pi$ to be linear in density up to first order, with $\chi$ the compressibility. Putting the above two equations together gives the equation for cell velocity:

$$\mu(\tau\partial_t v + v) - \mu v_0(\tau\partial_t p + p) = \eta\partial_x^2 v - \chi^{-1}\partial_x\rho \ . \tag{7}$$

Polarity dynamics is governed by the following equation,

$$\partial_t p = \kappa_p\partial_x^2 p + [a(\rho-\rho_c) - bp^2]p \ , \tag{8}$$

where the first term in the RHS induces alignment of polarity, while the second term causes breaking of symmetry in the homogeneous steady state when the density is above the critical threshold $\rho_c$. The strength of alignment and the degree of symmetry-breaking are controlled by the parameters $\kappa_p$ and $a$, respectively. We can non-dimensionalize Eqs. (4), (7) and (8) by defining rescaled variables as follows

$$t = \tau\bar{t}, \quad x = \sqrt{\frac{\eta}{\mu}}\bar{x} = \lambda\bar{x}, \quad \rho = \rho_0\bar{\rho}, \quad v = \frac{\lambda}{\tau}\bar{v}, \quad p = \frac{1}{\sqrt{\tau b}}\bar{p} \ . \tag{9}$$

These rescaled variables reduce the model equations to non-dimensional forms

$$\partial_{\bar{t}}\bar{\rho} = -\partial_{\bar{x}}(\bar{\rho}\bar{v}) + \bar{\kappa}\bar{\rho}(1-\bar{\rho}) \ , \tag{10.1}$$

$$\partial_{\bar{t}}\bar{v} = \partial_{\bar{x}}^2\bar{v} - \bar{v} - \bar{\chi}^{-1}\partial_{\bar{x}}\bar{\rho} + \bar{v}_0(\bar{p} + \partial_{\bar{t}}\bar{p}) \ , \tag{10.2}$$

$$\partial_{\bar{t}}\bar{p} = \bar{\kappa}_p\partial_{\bar{x}}^2\bar{p} + [\bar{a}(\bar{\rho}-\bar{\rho}_c) - \bar{p}^2]\bar{p} \ , \tag{10.3}$$

where

$$\bar{\kappa} = \kappa\tau, \quad \bar{\chi} = \frac{\eta\chi}{\rho_0\tau}, \quad \bar{v}_0 = \frac{v_0}{\lambda}\sqrt{\frac{\tau}{b}}, \quad \bar{\kappa}_p = \frac{\tau\kappa_p}{\lambda^2}, \quad \bar{a} = \rho_0\tau a, \quad \bar{\rho}_c = \frac{\rho_c}{\rho_0} \ . \tag{11}$$

#### B. Linear Stability Analysis

To determine the parameter regimes under which the local cell density remains bounded, we performed a linear stability analysis of the model equations about a homogeneous steady-state where polarity symmetry is broken. We linearized the fields as follows

$$\rho = \rho^*(t) + \delta\rho(x,t), \quad v = \delta v(x,t), \quad p = p^*(t) + \delta p(x,t) \ , \tag{12}$$

$$\partial_t\delta\rho = -\rho^*\partial_x\delta v + \bar{\kappa}(1-2\rho^*)\delta\rho \tag{13.1}$$

$$\partial_t\delta v = \partial_x^2\delta v - \delta v - \bar{\chi}^{-1}\partial_x\delta\rho + \bar{v}_0(\delta p + \partial_t\delta p) \tag{13.2}$$

$$\partial_t\delta p = \bar{\kappa}_p\partial_x^2\delta p + \bar{a}(\rho^* - \bar{\rho}_c - 3p^{*2})\delta p \tag{13.3}$$

We now perform a Fourier transform of the fields,

$$\mathcal{F}[\delta\rho] = \widetilde{\delta\rho} e^{i(qx-\omega t)}, \quad \mathcal{F}[\delta v] = \widetilde{\delta v} e^{i(qx-\omega t)}, \quad \mathcal{F}[\delta p] = \widetilde{\delta p} e^{i(qx-\omega t)} \tag{14}$$

which gives,

$$[\omega - i\bar{\kappa}(1-2\rho^*)]\widetilde{\delta\rho} - q\rho^*\widetilde{\delta v} = 0\,, \tag{15.1}$$

$$q\bar{\chi}^{-1}\widetilde{\delta\rho} - (\omega + i + iq^2)\widetilde{\delta v} + \bar{v}_0(\omega + i)\widetilde{\delta p} = 0\,, \tag{15.2}$$

$$[\omega + iq^2\bar{\kappa}_p - i\bar{a}(\rho^* - \bar{\rho}_c - 3p^{*2})]\widetilde{\delta p} = 0\,. \tag{15.3}$$

Solving for $\omega$ in terms of $q$ from

$$\begin{vmatrix} \omega - i\bar{\kappa}(1-2\rho^*) & -q\rho^* & 0 \\ q\bar{\chi}^{-1} & -(\omega + i + iq^2) & \bar{v}_0(\omega + i) \\ 0 & 0 & \omega + iq^2\bar{\kappa}_p - i\bar{a}(\rho^* - \bar{\rho}_c - 3p^{*2}) \end{vmatrix} = 0 \tag{16}$$

gives

$$\omega = i[\bar{a}(\rho^* - \bar{\rho}_c - 3p^{*2}) - \bar{\kappa}_p q^2],\, -\frac{i}{2}[1 + q^2 - \bar{\kappa}(1-2\rho^*)] \pm \frac{1}{2}\sqrt{4\bar{\chi}^{-1}q^2\rho^* - [1 + q^2 - \bar{\kappa}(1-2\rho^*)]^2} \tag{17}$$

In order for the density to be bounded from above, we need $\mathrm{Im}(\omega) < 0$ which gives the following conditions

$$\bar{a}(\rho^* - \bar{\rho}_c - 3p^{*2}) - \bar{\kappa}_p q^2 < 0 \tag{18}$$

and

$$\bar{\chi} < \frac{4q^2\rho^*}{[q^2 + 1 + \bar{\kappa}(1-2\rho^*)]^2}, \quad \bar{\kappa} < \frac{1+q^2}{1-2\rho^*} \tag{19.1}$$

Or

$$\frac{4q^2\rho^*}{[q^2 + 1 + \bar{\kappa}(1-2\rho^*)]^2} < \bar{\chi} < \frac{q^2\rho^*}{\bar{\kappa}(1-2\rho^*)(1+q^2)}\,. \tag{19.2}$$

Setting $q = \pi/2L$, $\rho^* = 1$, $p^* = \pm\sqrt{a(\rho^* - \bar{\rho}_c)}$ the conditions become

$$a < \frac{1}{6}\left(1 - \sqrt{1 - \frac{3\bar{\kappa}_p\pi^2}{L^2(1-\bar{\rho}_c)}}\right) \quad \text{or} \quad a > \frac{1}{6}\left(1 + \sqrt{1 - \frac{3\bar{\kappa}_p\pi^2}{L^2(1-\bar{\rho}_c)}}\right) \tag{20}$$

$$\bar{\chi} > \frac{\pi^2}{L^2}\left[\bar{\kappa} - \left(\frac{\pi^2}{4L^2} + 1\right)\right]^{-2} \tag{21}$$

where $L$ is half the length of the domain of solution.

#### C. Numerical implementation and parameter choices

We solved Eqs. (10) numerically through the finite volume approach using the FiPy module of Python. We assumed a domain $[-L, L]$ of size $2L$, much larger than tissue size, with no-flux boundary conditions for all variables, ie

$$\partial_x \rho = \partial_x v = \partial_x p = 0 \quad \text{at} \quad x = \pm L \, . \tag{22}$$

The initial density was taken to be a narrow Gaussian distribution centered at the origin,

$$\rho(x, t = 0) = \frac{1}{2} \exp\left(\frac{x^2}{2\sigma^2}\right) \tag{23}$$

where the standard deviation $\sigma$ (= 5 in our simulations) sets the initial size of the tissue. The initial polarity was randomly drawn from a normal distribution in the part of the domain within two standard deviations of the density profile, and zero elsewhere. The initial velocity was taken to be zero everywhere. The choice of model parameters for different simulation runs is summarized in Table II.

We determined the spatial extent of the tissue by stipulating that the density within the tissue is higher than a cutoff density of 0.01. Once we have determined the spatial extent of the tissue we can calculate its size, its midpoint and then the growth anisotropy index $g_{ans}$. During later stages of the simulation, the tissue starts collectively migrating in the direction of polarity and quickly reaches the boundary of the domain. To counter this behavior in the long simulation runs required for producing Fig. S8, we calculate the displacement of the tissue midpoint after every time step and displace all the variable profiles by an equal and opposite distance so the tissue remains centered at the origin. This does not affect the calculation of $g_{ans}$.

---

[7] R. Soerjadi. On the computation of the moments of a polygon, with some applications. 1968.

[8] Carsten Steger. On the calculation of arbitrary moments of polygons. 1996.

| Parameter | Symbol | Value | | | | | | |
|---|---|---|---|---|---|---|---|---|
| | | Fig. 1a-c | Fig. 2a-c | Fig. 2d | Fig. 2e | Fig. 3a-b | Fig. 3c | Fig. 3d |
| Initial cell number rate | $N_0$ | 1 | 9 | 9 | 9 | 9 | 9 | 9 |
| Proliferation rate | $G_0$ | 0.5 | 0.5 | 0.5 | 0.5 | 0.5 | 0.5 | 0.5 |
| Cell area elasticity | $K_A$ | 1 | 1 | 1-10 | 1 | 1 | 1-10 | 1 |
| Contact inhibition | $k$ | 100 | 100 | 100 | 10-100 | 100 | 100 | 10-100 |
| Variance of angular noise | $D_r$ | | | 1 | | | | |
| Edge tension | $\lambda$ | | | -0.02 | | | | |
| Motility | $v_0$ | 0.1 | 0.2 | 0-0.3 | 0.1 | 0.1 | 0-0.3 | 0-0.3 |
| Polarity alignment rate | $\kappa_p$ | 0 | 0.05 | 0.1 | 0.1 | 0.1 | 0.1 | 0.1 |
| Sizer | $A_S$ | | | 2 | | | | |
| Timer | $T$ | | | 5 | | | | |

TABLE I. Parameters for the vertex model simulations. The parameters are non-dimensional. In Fig. 3(a), the tissue without cell growth has $N_0 = 1001$ cells.

| Parameter | Symbol | Value | | |
|---|---|---|---|---|
| | | Fig. 4a | Fig. 4b-c | Fig. 4d |
| Proliferation rate | $\bar{\kappa}$ | 0.5 | 0.5 | 0.5 |
| Compressibility | $\bar{\chi}$ | 2 | 2 | 0.2 - 2 |
| Motility | $\bar{v}_0$ | 5 | 5 | 1 - 5 |
| Polarity alignment rate | $\bar{\kappa}_p$ | 0 | 1000 | 10 |
| Symmetry breaking coefficient | $\bar{a}$ | 0 | 0.3 | 0.35 |
| Threshold density | $\bar{\rho}_c$ | 0.2 | 0.2 | 0.2 |
| Half-length of simulation box | $L$ | 100 | 100 | 50 |

TABLE II. Parameters for the continuum model. The first six parameters are nondimensional.

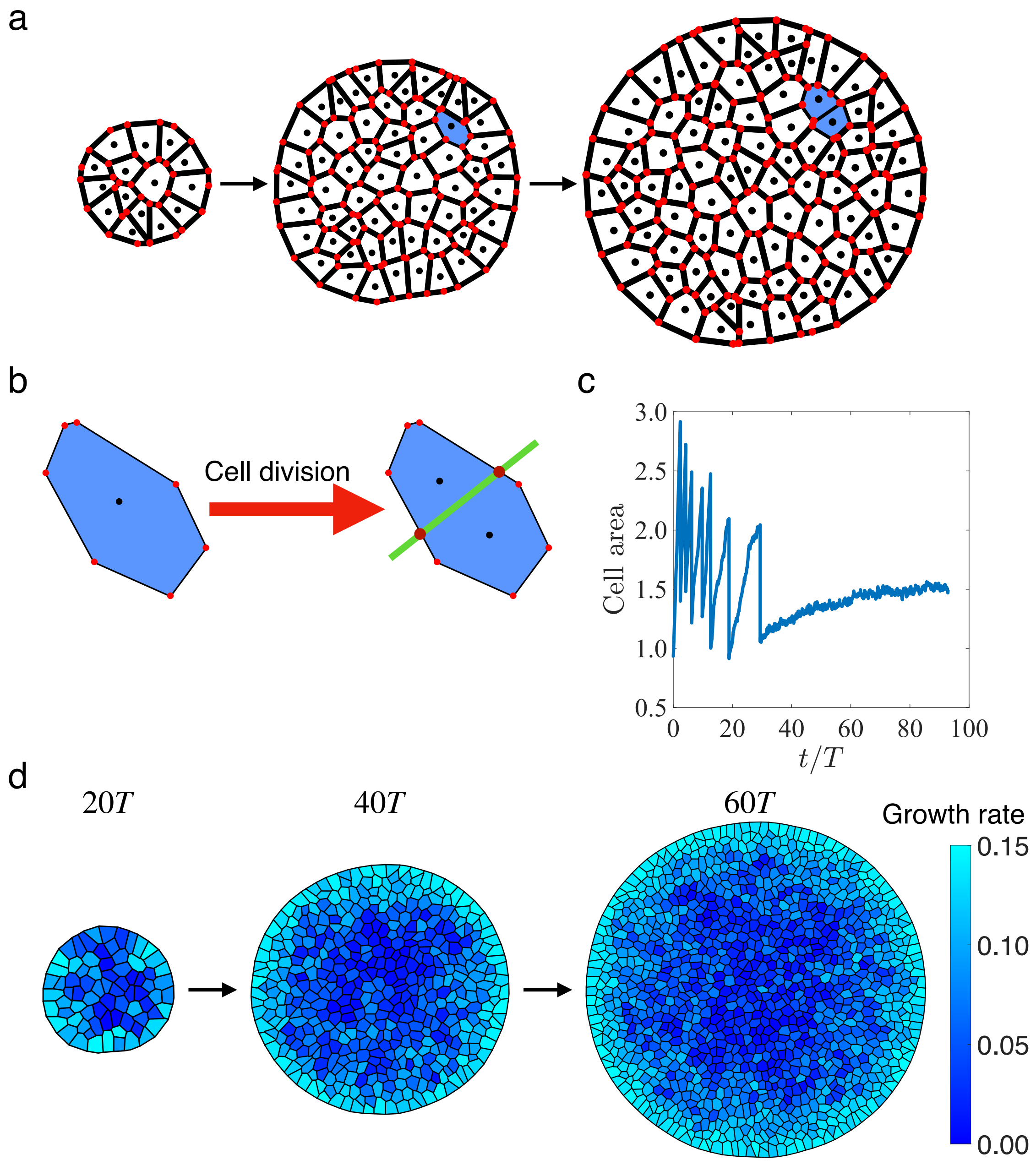


FIG. S1. **(a)** Time-lapse of configurations of an isotropically growing tissue. Black dots represent the cell centers and the red dots indicate the vertices obtained from voronoi tessellation. When the G1 sizer threshold is reached, a cell entering the G2 phase is marked by light blue in the snapshot. **(b)** Implementation of a cell division. When the cell reaches the timer threshold in the G2 phase, it divides in the direction perpendicular to the major axis (indicated by the green line). The two dark red dots represent the two new vertices connecting the new edge separating the daughter cells. **(c)** Single-cell area vs time for a representative cell in the bulk of the tissue, showing a transition to cell cycle arrest due to crowding-induced suppression of growth. **(d)** Timelapse of an isotropically growing tissue with random polarity. The colors represent cell growth rate which decreases from the boundary to the center of the tissue.

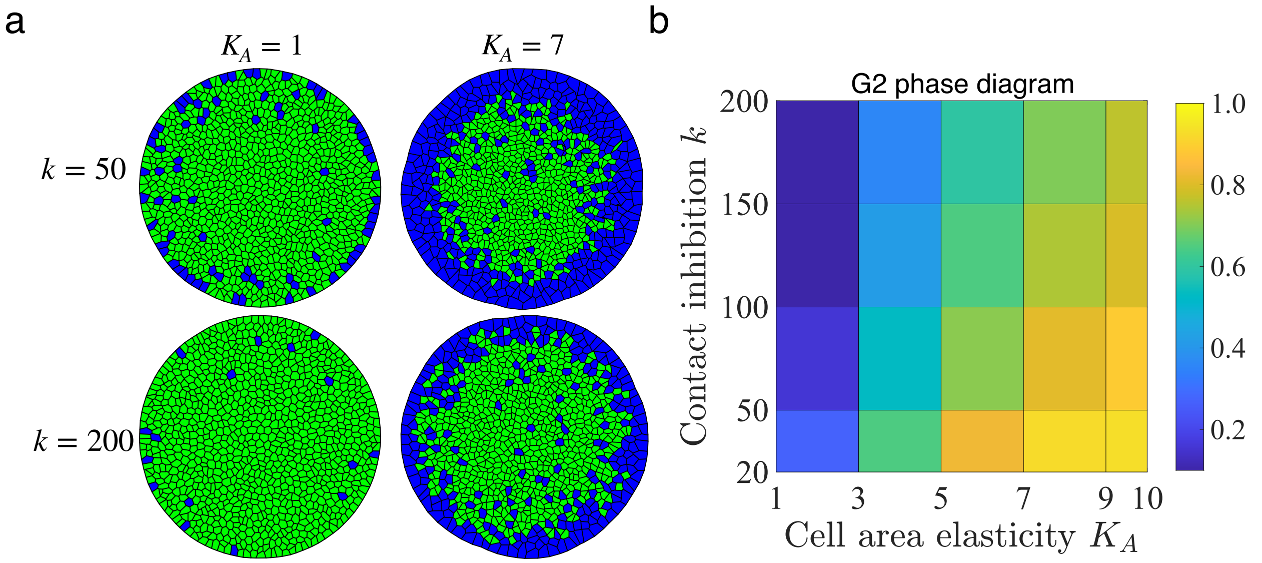


FIG. S2. **Patterns of cell cycle phase in a growing tissue by tuning contact inhibition $k$ and cell area elasticity $K_A$. (a)** Snapshots of growing tissues exhibiting boundary and bulk growth. Green cells are in the G1 phase and cells in the G2 phase are colored in blue. **(b)** Phase diagram of the ratio of cells in G2 phase. At a fixed contact inhibition $k$, increasing $K_A$ leads to more fraction of cells in the G2 phase. In contrast, increasing $k$ at a given $K_A$ value, cells are more sensitive to crowding and fewer cell divisions could occur in the tissue bulk, resulting in a lower fraction of cells in the G2 phase.

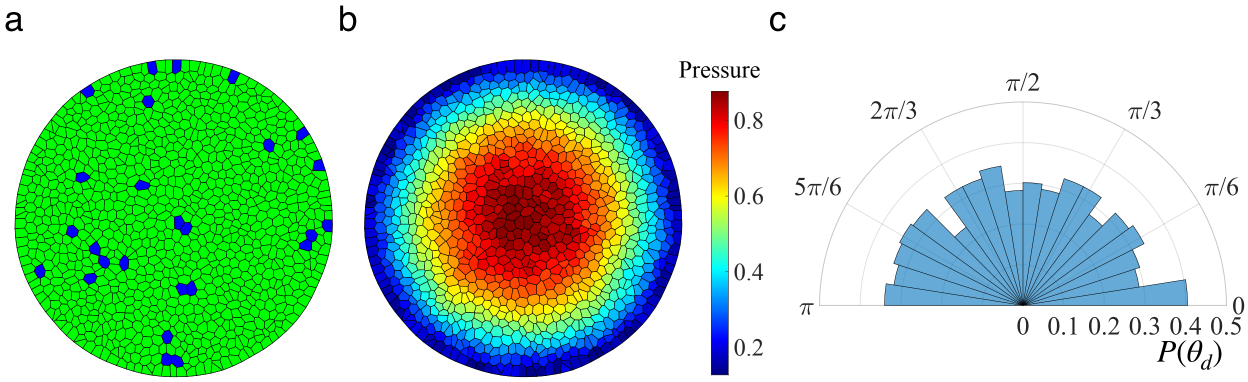


FIG. S3. **Growth pattern in the absence of contact inhibition.** Spatial map of **(a)** cells in G1 (green) and G2 (blue) phases and **(b)** pressure distribution for a growing tissue with $k = 0$. **(c)** The angular distribution of cell divisions shows isotropy.

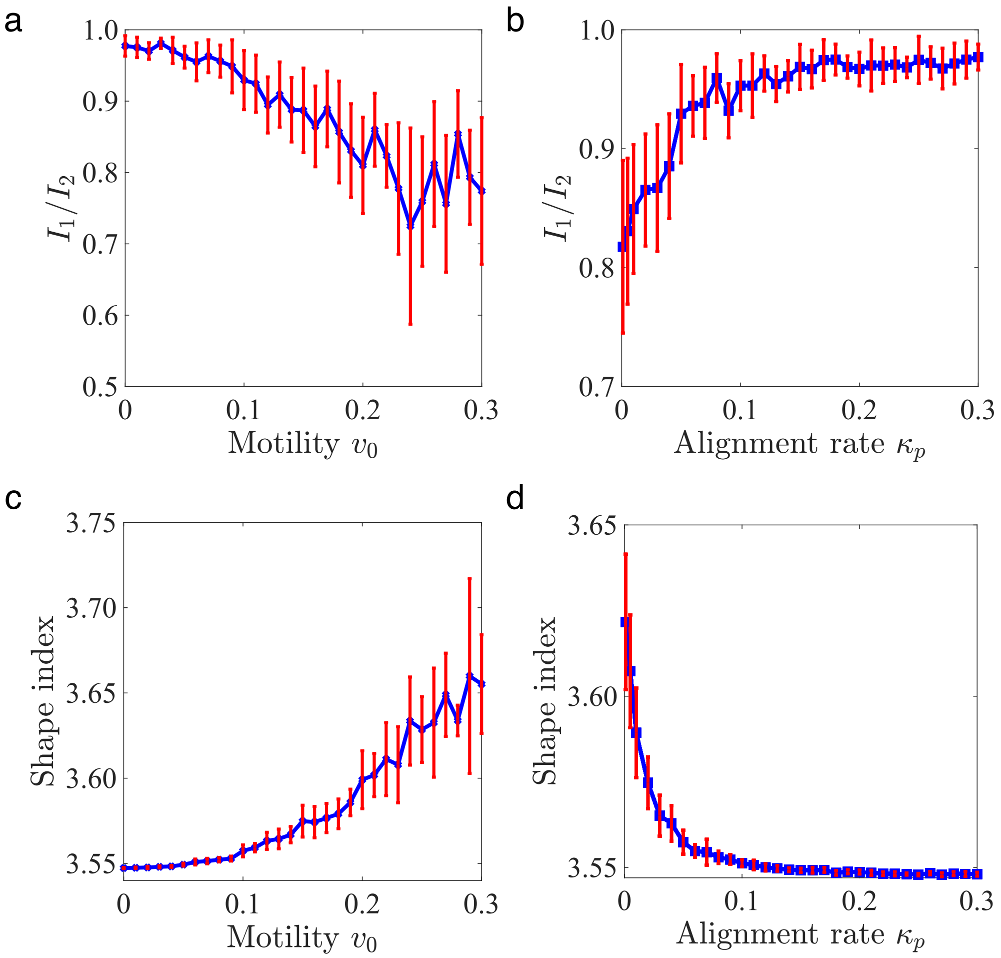


FIG. S4. **Control of tissue shapes by cell motility and polarity alignment rate.** At $\kappa_p = 0.05$, we characterize the tissue aspect ratio **(a)** and tissue shape index **(c)** as a function of cell motility $v_0$. With the increase of $v_0$, the cell division anisotropy increases and tissue shapes become non-round like. At $v_0 = 0.1$, the tissue aspect ratio **(b)** and tissue shape index in **(d)** is plotted as a function of the polarity alignment rate $\kappa_p$. When $\kappa_p$ is increasing, the polarity vectors are more aligned, leading to the collective drifting of the tissue. The tissue shapes are more round-like.

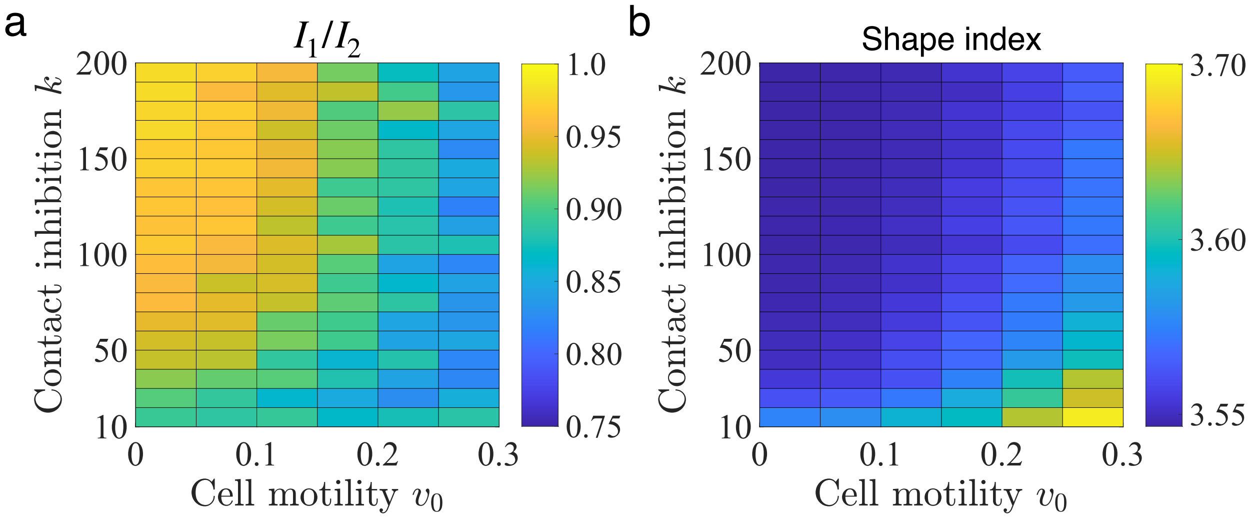


FIG. S5. **Characterization of tissue shapes at different values of cell motility and contact inhibition.** Phase diagram of tissue aspect ratio **(a)** and shape index **(b)** by tuning contact inhibition $k$ and cell motility $v_0$. $k$ suppresses while $v_0$ promotes the anisotropic tissue shapes.

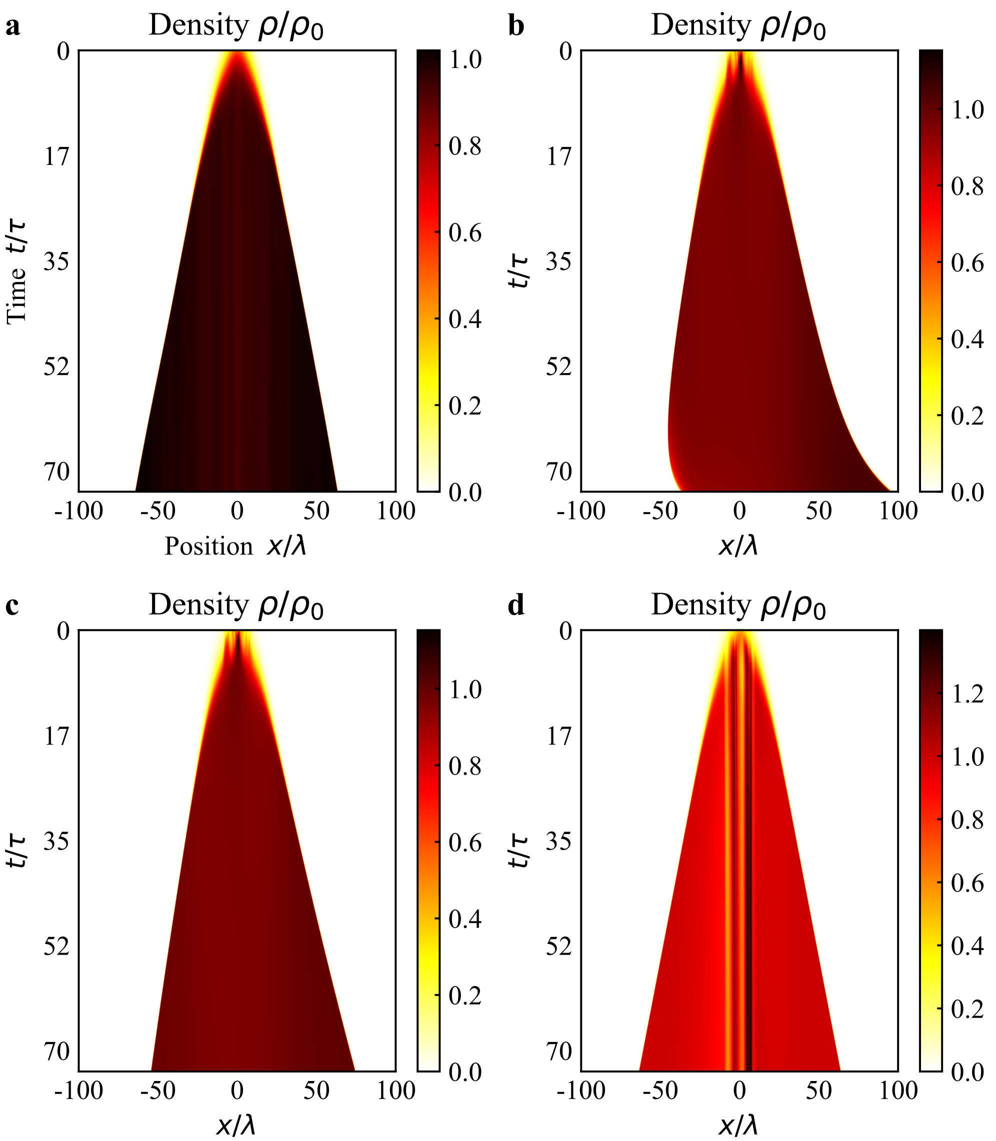


FIG. S6. **Effects of cell polarity on the pattern of growth.** When $\kappa_p = 0, a = 0$ **(a)** and polarity is randomised for every time step, growth is isotropic. When both $\kappa_p$ and $a$ are nonzero ($\kappa_p = 1000$, $a = 0.3$ **(b)**) the tissue grows anisotropically. For $\kappa_p = 1000$, $a = 0$ **(c)** the anisotropy is much less pronounced. In the presence of only the symmetry breaking term ($\kappa_p = 0$, $a = 0.3$ **(d)**) the initial fluctuations in the density, due to the initial random configuration of polarity, persist through time.

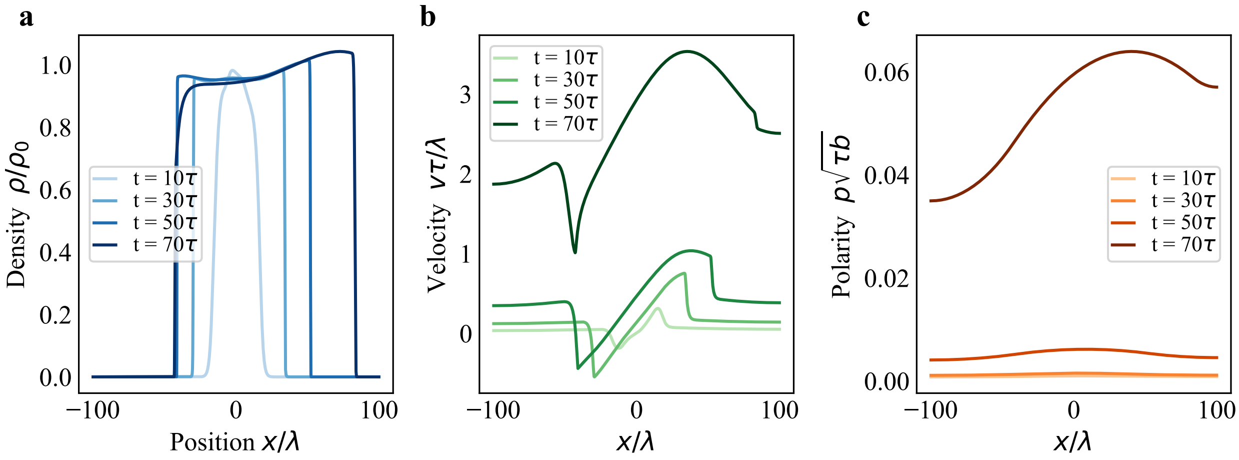


FIG. S7. **Spatial profiles of density, polarity and velocity during anisotropic growth.** Snapshots of **(a)** Density, **(b)** Velocity and **(c)** Polarity during different times for the anisotropically growing tissue shown in Fig. 4b and Fig. S6b.

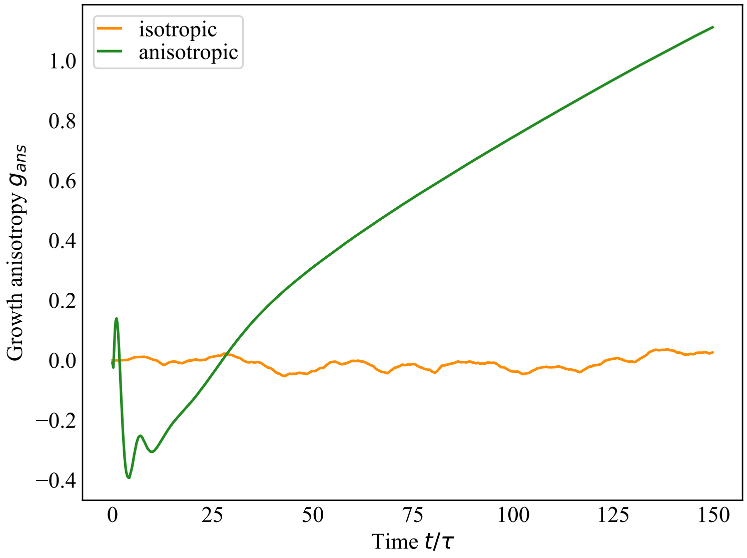


FIG. S8. **Growth anisotropy index for isotropic and anisotropic growth.** The growth anisotropy index $g_{ans} \approx 0$ throughout the simulation for isotropically growing tissue, but keeps increasing for the anisotropic case after some initial fluctuations.
